# Bridging Organ Transcriptomics for Advancing Multiple Organ Toxicity Assessment with a Generative AI Approach

**DOI:** 10.1101/2024.04.02.587739

**Authors:** Ting Li, Xi Chen, Weida Tong

**Author notes:** **Corresponding author:** Weida Tong.

## Abstract

Translational research in toxicology has significantly benefited from transcriptomic profiling, particularly in drug safety. However, its application has predominantly focused on limited organs, notably the liver, due to resource constraints. This paper presents TransTox, an innovative AI model using a Generative Adversarial Network (GAN) method to facilitate bidirectional translation of transcriptomic profiles between the liver and kidney under drug treatment. TransTox demonstrates robust performance, validated across independent datasets and laboratories. Firstly, the concordance between real experimental data and synthetic data generated by TransTox was demonstrated in characterizing toxicity mechanisms compared to real experimental settings. Secondly, TransTox proved valuable in gene expression predictive models, where synthetic data could be used to develop gene expression predictive models or serve as “digital twins” for diagnostic applications. The TransTox approach holds potential for multi-organ toxicity assessment with AI and advancing the field of precision toxicology.

## Introduction

Translational science plays a pivotal role in various domains. This field encompasses a wide array of translational research focus with practical applications. For instance, transitioning genomics data between platforms, such as from microarray to next-generation sequencing, offers a way to capitalize on prior investments ^1^. Moreover, translating data across diverse biological contexts, such as from non-invasive blood samples to tissue samples, holds significant promise for clinical practice ^2^. In toxicology, it facilitates cross-assay extrapolation, notably the translation of in vitro findings to in vivo observations, known as in vitro to in vivo extrapolation (IVIVE). For drug safety, it bridges the gap between preclinical animal investigations and clinical outcomes. Recent advancements in artificial intelligence (AI), particularly in generative AI, have offered novel ways to augment translational research, exemplified by techniques like Generative Adversarial Networks (GANs) and diffusion models. In this study, we introduce an AI-driven approach capable of predicting toxic effects in one organ based on observations from another, thus bolstering systems toxicology.

Animal models play a crucial role in evaluating drug safety. The analysis of gene expression profiles from animal studies, a widely employed toxicogenomics (TGx) approach, has significantly contributed to the drug safety assessment in two significant ways. Firstly, it elucidates the underlying biological processes of toxicity in animals and their relevance to humans ^3–9^. Secondly, it facilitates the development of gene expression-based predictive models for toxicant classification ^10–18^. Given the resource-intensive nature of conducting comprehensive TGx studies for every organ, most TGx studies have been centered on a limited number of organs, where the liver TGx experiment is dominant ^19^. However, multiple studies have demonstrated that toxicity could manifest across multiple organs under chemical exposure or drug treatment ^20,21^. For example, doxorubicin, used as an anti-neoplastic drug in the treatment of various types of cancers, causes heart, liver, and kidney injury ^22^. While it’s important to study individual organs, it’s equally crucial to consider the interactions and interdependencies between different organs and systems within an organism. Consequently, toxicity assessments often involve evaluating the effects on multiple organs to gain a comprehensive understanding of the overall impact on the organism’s health ^23,24^. Particularly, advancements in systems biology and computational modeling have been focused on exploring the complex interactions between organs and predict potential toxic effects more accurately at an organism level ^25–28^.

Generative Adversarial Networks (GANs) have gained prominence in biomedicine for applications such as generating new molecules ^29–32^. In toxicology, GANs have been used to generate synthetic data that closely mimic the study results from real biological samples ^10,33–36^. Considering the intricate interactions and interdependencies among various organs, we developed a GAN-based model to explore the possibility of translating gene expression profiles between organs of healthy rats ^37^. This proof-of-concept study laid the foundation to investigate the transcriptomic translation between different organs under the treatment conditions. Different organs, intricately connected through various feedback mechanisms, likely respond to drug treatment in an interconnected fashion. Such cross-organ translation not only broadens the scope of toxicological investigation but also improves a holistic understanding of the systemic impact of treatments and substances under investigation.

The liver and kidneys are primary target organs for toxicity^38^ due to their shared roles in detoxification and metabolism^39^. They share important toxicity signaling pathways involving oxidative stress, inflammation, and apoptosis^39–41^. Damage to one organ can affect the other through systemic inflammation and altered signaling molecules^42–46^. Additionally, they both express drug metabolizing enzymes and transporters (DMETs), such as cytochrome P450 enzymes (CYPs)^47^, solute carrier (SLC), and ATP-binding cassette (ABC) transporters^48–50^. This cross-organ interdependence suggests the potential of translating findings between them, where the cumulative burden of toxic substances on both organs may reflect not only in their biological functions but also in their molecular expression. Particularly, given the fact that most TGx studies have been focused only on the liver, this study may offer an opportunity to infer transcriptomic responses of kidney based on the historical liver data.

In this study, we developed TransTox to enable the bidirectional translation of transcriptomic profiles between the liver and kidney under drug treatment. TransTox was trained using transcriptomic data from the Toxicogenomics Project-Genomics Assisted Toxicity Evaluation Systems (TG-GATEs) ^51^, which included varying doses (high/medium/low/control) and treatment durations (3/7/14/28 days). The model was evaluated using three metrics —cosine similarity, root mean square error (RMSE), mean absolute percentage error (MAPE)—and one visualization approach—Uniform Manifold Approximation and Projection for Dimension Reduction (UMAP) ^52^. Importantly, the model was assessed on independent datasets from both the training set’s source labs and a different lab. To evaluate the synthetic data generated by TransTox in elucidating toxicity mechanisms, we compared the synthetic differential expressed genes (DEGs) and enriched pathways with those obtained from real experimental settings. Additionally, we evaluated the value of TransTox in gene expression predictive models, where synthetic data could be used to develop valid gene expression predictive models or serve as “digital twins” for diagnostic applications. This comprehensive study demonstrates the potential of TransTox in translating transcriptomic profiles between organs to advance TGx research.

## Results

### TransTox Training Results

The TransTox model comprises two sets of generator and discriminator pairs, one for the liver and the other for the kidney. These components interact as shown in **Figure 1**. The model was developed using the training set consisted of 45,402 pairs of liver and kidney profiles, corresponding to 32 compounds (approximately 80% of the total 41 compounds) under 510 treatment conditions, involving 1,526 rats. The liver and kidney profiles were obtained from the TG-GATEs database ^51^ (see Methods). Both liver and kidney generators were selected at 1,000 epochs, where the loss function was stabilized. High agreement was observed between synthetic and real profiles for both the liver and kidney, as visualized with UMAP (**Figure 2a, e**). To objectively assess the training results, we included a negative control based on the real (experimental) data. The negative control was used to establish a baseline for assessing the predictive accuracy of gene expression in one organ based on profiles from a different organ. In the negative control group, measurements such as cosine similarity were calculated between any two real transcriptomic profiles (excluding biological duplicates) within an organ. In the liver, the agreement between synthetic and real data was 0.9995 in cosine similarity, significantly higher than the mean of the negative control group (0.9980) with a t-test p-value <0.001 (**Figure 2b**). Similarly, there was a small RMSE of 0.247, significantly lower than the mean of the negative control group (0.488) with a p-value <0.001 (**Figure 2c**). The MAPE was 0.025, also significantly smaller than the negative control group’s mean of 0.050 with a p-value <0.001 (**Figure 2d**). Similar results were obtained in kidney, with a high cosine similarity of 0.9996 (against the negative control of 0.9986 with a p-value <0.001, **Figure 2f**), RMSE of 0.218 (compared to the negative control of 0.419 with a p-value <0.001, **Figure 2g**), and MAPE of 0.019 (against the mean of the negative control of 0.039 with a p-value <0.001, **Figure 2h**). These results suggest that the high agreement between the synthetic and real profiles for both liver and kidney.

**Figure 1:**
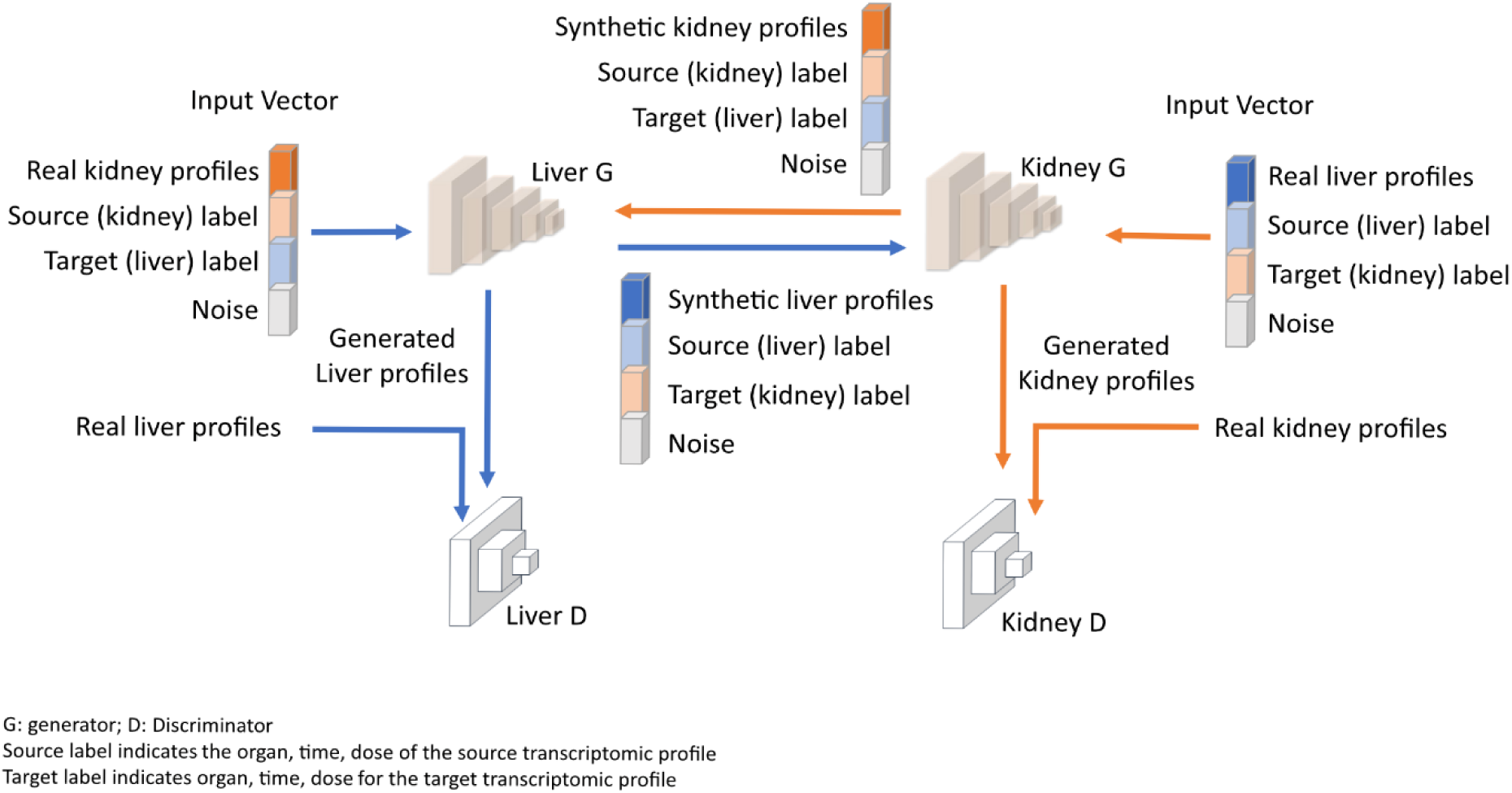
TransTox Framework. The architecture includes a liver generator and a kidney generator, each paired with a liver discriminator and a kidney discriminator. The generator’s input comprises four components: the real transcriptomic profile from the source organ, label information for the source organ, label information for the target organ, and a noise vector. **G**: generator; **D**: Discriminator; Source label indicates the organ, time, dose of the source (initial) transcriptomic profile (like kidney, 3-day, low); Target label indicates organ, time, dose for the target transcriptomic profile (like liver, 28-day, middle).

**Figure 2:**
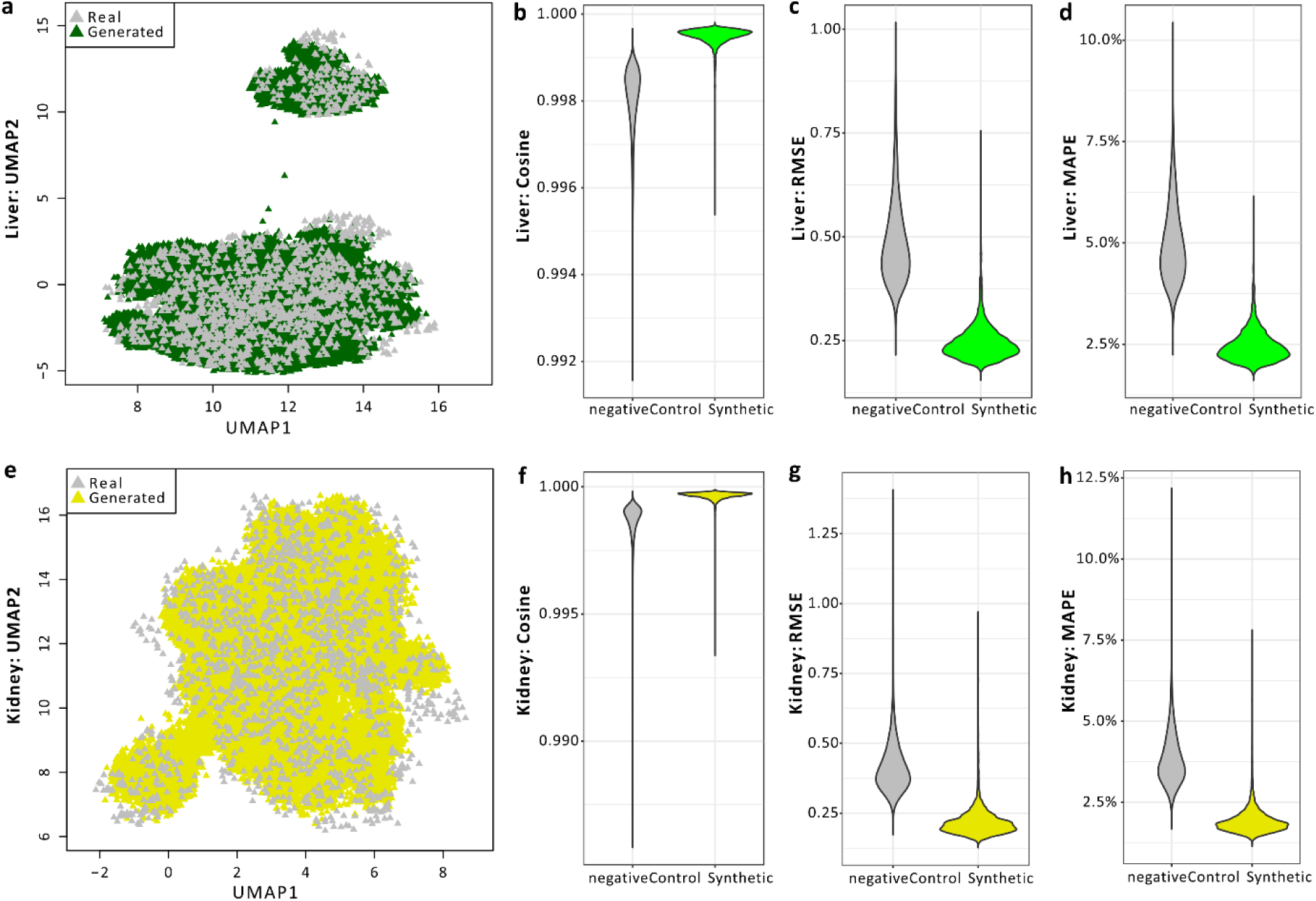
Training performance of TransTox. Evaluation using UMAP, cosine similarity, RMSE, and MAPE for liver (**a** to **d**) and kidney (**e** to **h**). In negative control group, measurements such as cosine similarity were calculated between any two real transcriptomic profiles (excluding biological duplicates) within an organ. In synthetic group, each measurement was calculated between the synthetic profiles and its corresponding real profiles.

### Validation of TransTox Predictions with a Test Set from the Same Lab

The TransTox model was evaluated with a TG-GATEs test set that consisted of 11,250 pairs of profiles, corresponding to 9 compounds (approximately 20% of the total 41 compounds) under 106 treatment conditions, involving 318 rats. We employed the same four approaches as the training set to evaluate the test results. For both the liver and kidney, the predicted profiles showed a high degree of agreement with the experimental results, as illustrated in the UMAP visualization and quantitively measured by cosine similarity, RMSE, and MAPE (**Supplementary Figure 1**).

Furthermore, we conducted a comparison at the gene level, examining the treatment-based synthetic profiles alongside their corresponding real profiles for each organ and between organs. We first calculated the difference in gene expression values between the synthetic and real data (normalized to the real value) for the liver, presented in a histogram plot. As depicted in **Figure 3a**, 95% of synthetic gene expression values fall within a range of less than 10% variation compared with real values. We then conducted the same analysis for the kidney, where 98% of synthetic gene expression values were within less than 10% variation in comparison to real values (**Figure 3b**). Lastly, to ensure the relative expression between two organs for each gene remained unchanged, we calculated the difference between synthetic and real data with respect to the between-organ gene distance (normalized to the average of the real value of the two organs) for all treatment conditions. As shown in **Figure 3c**, the majority (93%) of the synthetic between-organ gene distances were maintained with variations of less than 10% in comparison to real between-organ gene distances.

**Figure 3:**
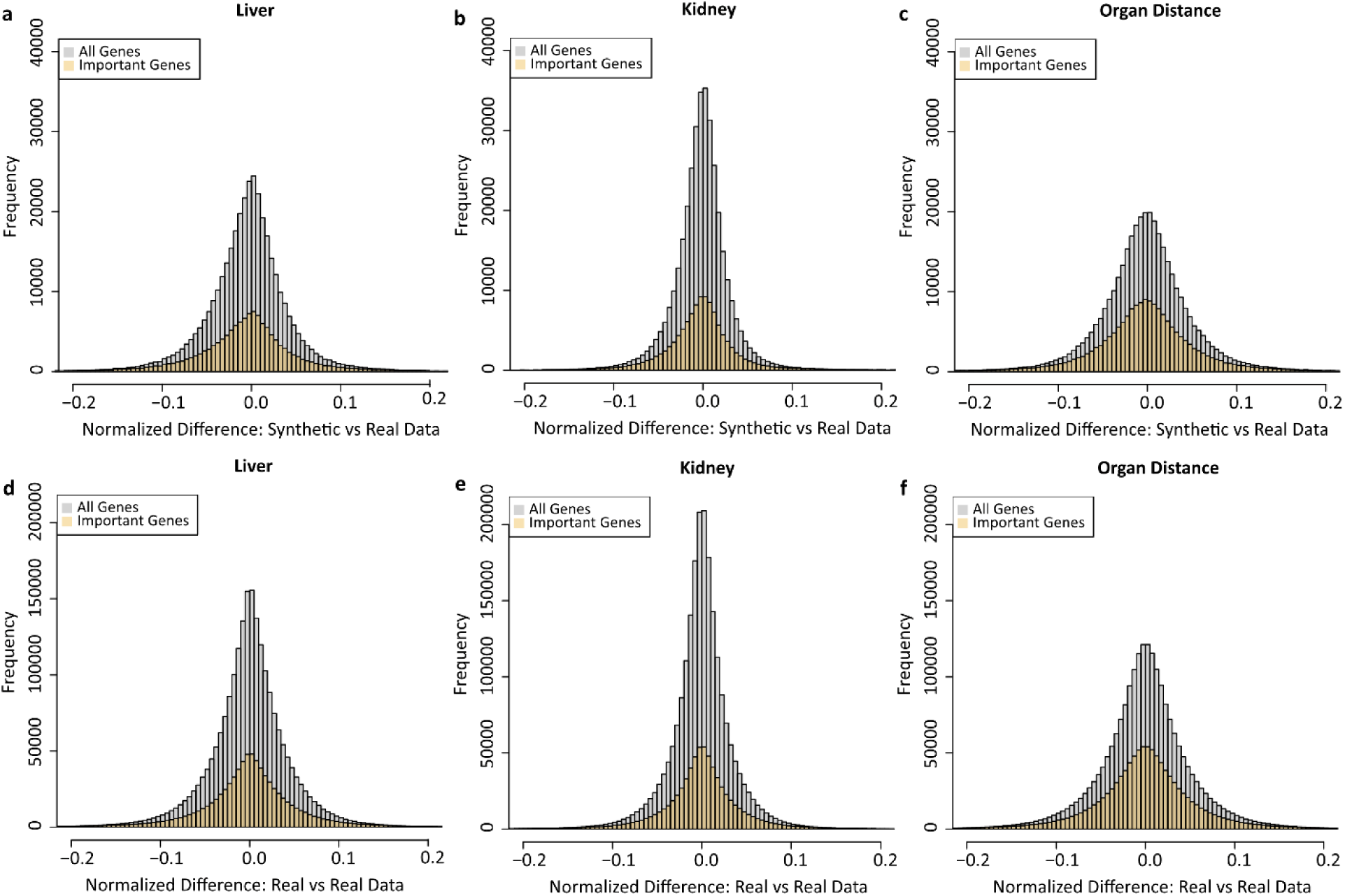
Gene-level comparison of TransTox in the TG-GATEs test set. The comparison is conducted at the gene level between synthetic and real treatment-based profiles, both within and between organs. The x-axis represents normalized differences, while the y-axis shows the number of gene-treatment combinations. **a** Difference between synthetic and real liver profiles. **b** Difference between synthetic and real kidney profiles. **c** Between-organ gene differences: synthetic profiles vs. real profiles. In addition, we also provide the gene-level comparison between the biological duplicates, **d**, **e**, and **f** for liver, kidney, and between-organ differences, respectively. “All genes” refer to the studied S1500 genes, while “Important Genes” are those with an absolute log2-transformed fold change greater than 1 in at least one treatment.

Given that not all genes are equally important with respect to mechanistic study and biomarker development, we defined a subset of genes, called Important Genes, which had an absolute log2-transformed fold change greater than 1 in at least one treatment condition (**Supplementary Data 1**). As shown in **Figure 3**, similar trends were observed for these genes.

If we consider TransTox as a pseudo-experiment, its synthetic results need to be compared with the variability observed in real experiment. For that, we calculate variability in transcriptomic data based on differences in gene expression values among biological duplicates. As shown in the **Figure 3d**, the variability in liver among biological duplicates exhibited a distribution similar to the differences between synthetic and real values in the liver (**Figure 3a**). Specifically, 95% of gene expression values among biological duplicates varied by less than 10% (**Figure 3d**). We performed the same analysis for the kidney (**Figure 3e**) and organ distance (**Figure 3f**) among biological duplicates. We found that the pattern of differences between synthetic and real gene expression values was consistent with the pattern observed among biological duplicates.

### Analysis of Organ-Specific Genes and Mechanisms with Thioacetamide Treatment

Some compounds cause toxicity to the liver but not to the kidneys. The question arises: how can changes in liver-specific genes caused by these compounds predict the expression of genes specific to the kidneys, which should not be affected by liver changes? To test this, we analyzed four organ-specific genes under treatment with thioacetamide, a compound known to cause liver toxicity with minimal impact on the kidneys. This analysis aimed to further evaluate the reliability of TransTox predictions by assessing the agreement between synthetic and real data for these organ-specific genes treated with organ-specific toxicants. Specifically, we analyzed four organ-specific genes: two predominantly expressed in the liver and two in the kidneys. For the liver, we examined cholesterol 7 alpha-hydroxylase (*Cyp7a1*) and Alpha-fetoprotein (*Afp*). *Cyp7a1* is primarily expressed in the endoplasmic reticulum of hepatocytes^53^, while AFP is commonly used as a biomarker for hepatocellular carcinoma surveillance and plays a role in liver regeneration^54^. For the kidneys, we analyzed aquaporin 2 (*Aqp2*) and paired box 8 (*Pax8*). AQP2 is exclusively expressed in the principal cells of the connecting tubule and collecting duct and is the predominant vasopressin-regulated water channel^55,56^, while *Pax8* is a central regulator of kidney development^57^. As shown in **Figure 4**, the mean values of these four genes were comparable between the synthetic and real data, except for *Afp*, for which the synthetic data was slightly lower than the real expression value.

**Figure 4:**
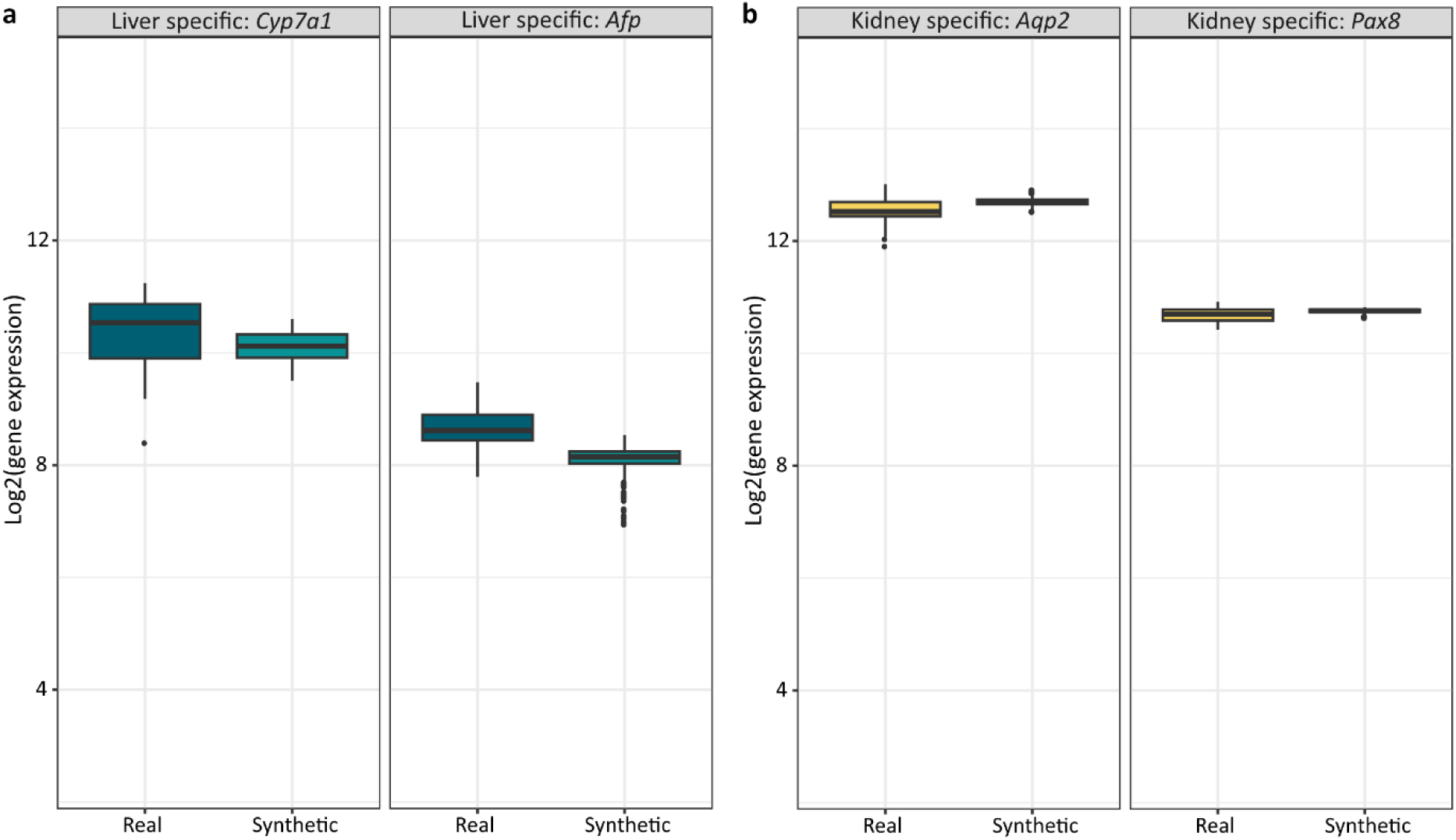
Organ specific gene expression comparison. The comparison is between real and synthetic profiles under Thioacetamide treatments. **a** Liver specific genes: *Cyp7a1* and *Afp*. **b** Kidney specific genes: *Aqp2* and *Pax8*.

We also evaluated the synthetic liver profiles predicted from kidney data under high-dose, 28-day thioacetamide treatment, which represents a challenging translation due to severe liver injury but minimal kidney impact in rats. The synthetic profiles identified differential expression of key toxicity-related genes, including *Cyp2b1*, *Cyp7a1*, and *Cyp2b2* from the cytochrome P450 family, known for their roles in thioacetamide toxicity after metabolic activation^58^. Additionally, KEGG pathway analysis of differentially expressed genes from the synthetic profiles revealed enrichment in the “Metabolism of Xenobiotics by Cytochrome P450” pathway, consistent with the findings above. The enrichment analysis also identified other relevant pathways such as ABC transporters, AGE-RAGE signaling in diabetic complications, bile secretion, chemical carcinogenesis, cholesterol metabolism, drug metabolism, PPAR signaling, steroid biosynthesis, steroid hormone biosynthesis, and ketone body synthesis and degradation. Some of these pathways are reported to be important in thioacetamide mechanisms^59,60^.

### External Validation of TransTox Predictions Using Data from a Different Lab

We evaluated TransTox performance using data from the DrugMatrix dataset (**Supplementary Data 2**), which originated from a different laboratory with a similar but not identical study design (see Methods section). The generated profiles were compared with the actual profiles in the DrugMatrix using cosine similarity (**Figure 5**).

**Figure 5:**
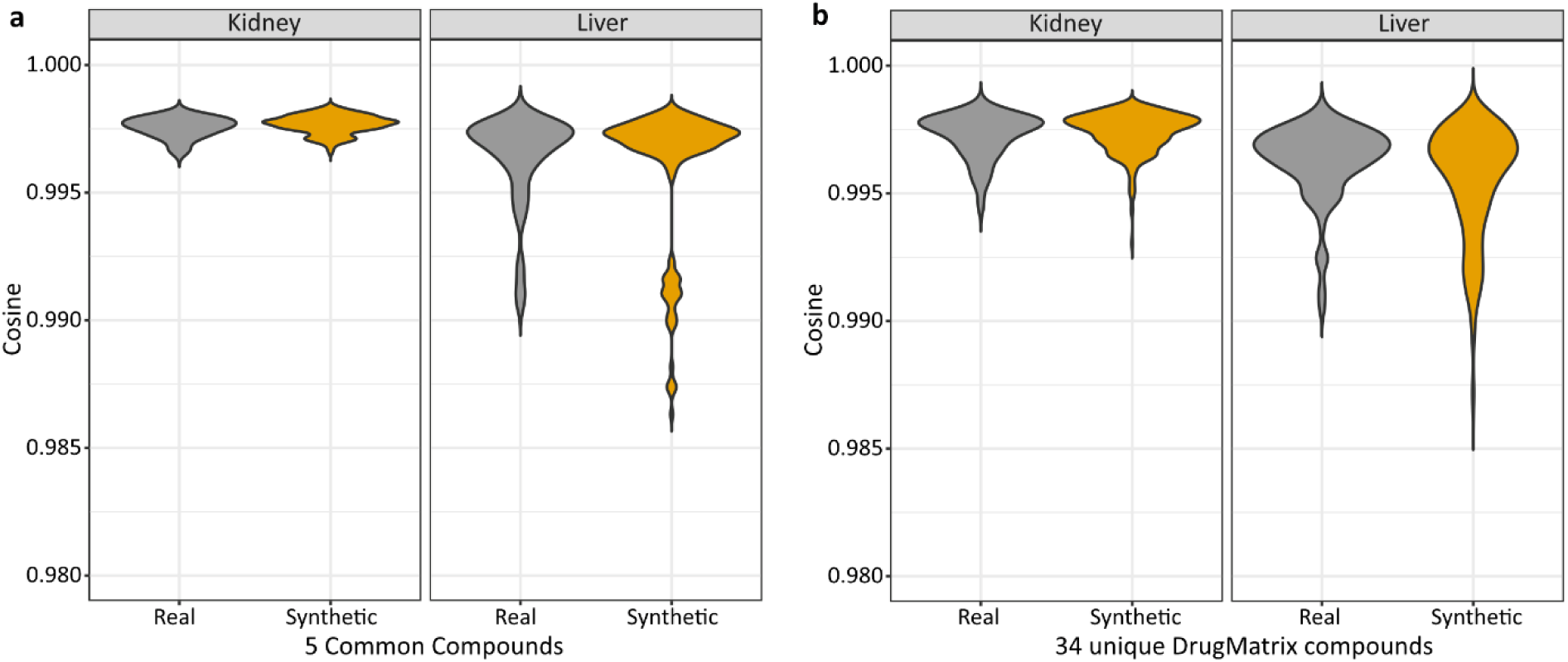
TransTox’s performance on the external validation set of DrugMatrix. a The cosine similarity distribution is shown for the five common compounds between DrugMatrix and TG-GATEs. The “Real group” represents values calculated between the real DrugMatrix and TG-GATEs transcriptomic profiles, while the “Generated group” indicates values calculated between the real DrugMatrix and synthetic transcriptomic profiles generated by TransTox. b The cosine similarity distribution is presented for the 34 compounds for which DrugMatrix had both liver and kidney profiles, but TG-GATEs had data available for only one organ or neither.

There were five compounds for which both liver and kidney profiles were available and matched between TG-GATEs and DrugMatrix. This provided an opportunity to assess the accuracy of synthetic data compared to the differences observed between the two labs using real experimental data. **Figure 5a** comprises four violin plots: two for the kidney (left panel) and two for the liver (right panel). For each organ, the left violin plot represents the correlation of experimental data, while the right box plot represents the correlation between synthetic and real profiles. For both organs, no statistical difference was observed in the correlation between real and generated profiles compared to the correlation of real profiles between TG-GATEs and DrugMatrix.

There were 34 compounds from DrugMatrix that had both liver and kidney profiles, but they had incomplete or no data in TG-GATEs. TransTox was applied to generate synthetic profiles for both the liver and kidney, as shown in the right violin plots of each panel in **Figure 5b**. Similar to **Figure 5a**, we employed two reference sets (one for the liver and the other for the kidney) to evaluate the accuracy of the synthetic results. For the liver, 29 compounds had profiles available in both TG-GATEs and DrugMatrix, with 9 compounds overlapping the 34 compounds. Therefore, we used the liver profiles of these 29 compounds as a reference point. However, for the kidney, only 11 compounds had profiles available in both datasets, and none overlapped with the 34 compounds. In summary, the similarity of the real and generated data was slightly lower (0.9955) than that of the reference data (0.9963) in the liver, while comparable for the kidney where the similarity of the generated group and real group (reference data) was 0.9974 and 0.9973, respectively.

### TransTox-Generated Transcriptomic Profiles in Elucidating Toxicity Mechanisms in Comparison with Real *Experimental* Settings

To assess the ability of TransTox to recapitulate results from the real TGx studies, especially in terms of mechanistic elucidation, we compared synthetic profiles with their corresponding real profiles, focusing on DEGs and enriched pathways (see Methods section). The comparison was conducted using the test set.

**Figure 6a** depicted the percentage of overlapped DEGs between synthetic and real profiles across three dose levels and four time points in the liver test set. Variations were observed for different dose and time points. The average overlapped DEGs ratios were 0.426, 0.436, and 0.364 for the low, middle, and high dose groups, respectively, and ratios of 0.410, 0.396, 0.381, and 0.454 for the 3-, 7-, 14-, and 28-days treatments, respectively. When comparing the percentage of overlapped KEGG pathways between two sets of DEGs of synthetic and real profiles, the ratio of overlapped pathways (with p-value < 0.05) as presented in **Figure 6b** was 0.349, 0.422, and 0.437 for the low, middle, and high dose groups, respectively, and 0.297, 0.435, 0.389, and 0.488 for the 3-, 7-, 14-, and 28-days treatments, respectively.

**Figure 6:**
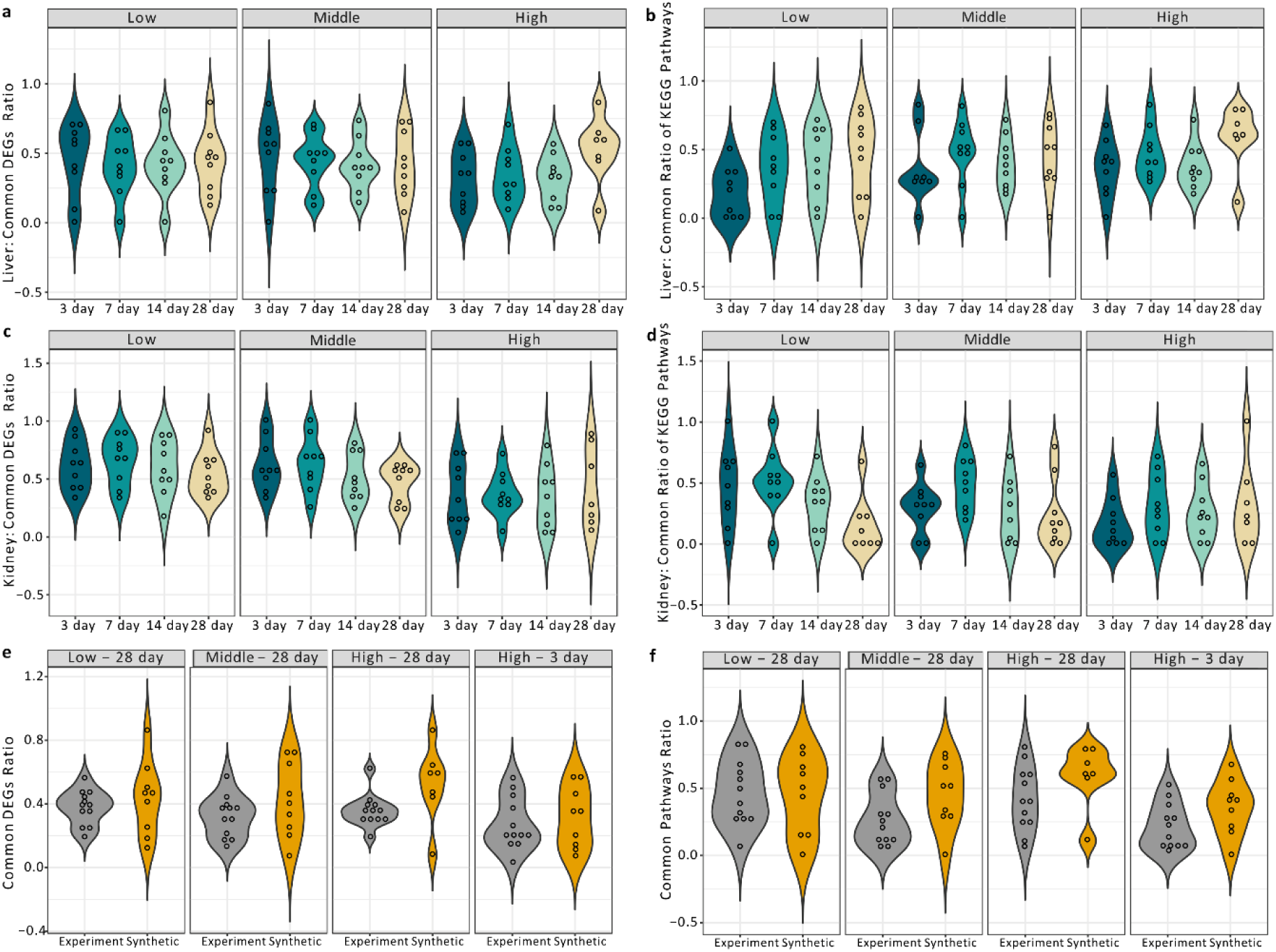
Comparison of differentially expressed genes (DEGs) and pathway enrichment analyses. DEGs identified by synthetic profiles were compared to those defined by real transcriptomic profiles. DEGs from both synthetic and real profiles were independently subjected to KEGG pathway enrichment analyses. The Violin plots illustrate the overlap ratio of DEGs and enriched KEGG pathways in comparison to real profiles under different dose and time conditions. Liver results are depicted in panels **a** and **b**, while kidney results are presented in panels **c** and **d**. Panels **e** and **f** illustrate the overlap ratio of DEGs and pathways for real experimental settings subjected to the same treatments for two compounds, Entacapone and Tolcapone, while the generated group presents the overlap ratio of synthetic and real liver profiles under the corresponding treatment condition.

Similar analyses were performed for synthetic profiles in the kidney test set, and variations were also observed among different dose and time points. The average overlapped DEGs ratios were 0.597, 0.553, and 0.368 for the low, middle, and high dose groups, respectively, and 0.529, 0.547, 0.478, and 0.478 for the 3-, 7-, 14-, and 28-days treatments, respectively (**Figure 6c**). Additionally, KEGG pathway enrichment analyses yielded an average overlapped ratio of enriched KEGG pathways (with a p-value < 0.05) of 0.369, 0.323, and 0.258 for low, middle, and high dose groups, respectively, and 0.306, 0.435, 0.286, and 0.231 for the 3-, 7-, 14-, and 28-days treatments, respectively (**Figure 6d**).

To contextualize the above results within real experimental settings, we selected a TGx study involving two drugs (entacapone and tolcapone) with a similar study design and transcriptomics experiment as TG-GATEs. This study used 12 rats for each treatment group, allowing them to be divided into four random groups of 3 rats each, aligning with TG-GATEs. We assumed that each group represented an independent experiment, and the identical experiment was conducted four times, with the only difference being the rats used (see Methods section). Subsequently, we analyzed the overlap in DEGs and pathways between any two groups to serve as reference points for evaluating the performance of TransTox results. As presented in **Figure 6e**, the overlap in DEGs between two identical experiments was in a similar range as observed in the comparison between synthetic and real profiles presented in **Figure 6a**. Similarly, the overlap in enriched KEGG pathways for any two identical experiments relating to entacapone and tolcapone (**Figure 6f**) was also in a similar range as observed in the comparison between synthetic and real profiles depicted in **Figure 6b**.

### TransTox for Gene Expression Predictive Model Development – Necrosis Prediction

To assess the potential of TransTox in gene expression predictive model development, we explored its application in identifying necrosis, a prevalent pathological outcome in both drug-induced liver and kidney injuries. Necrosis is a characteristic feature of acute and chronic liver diseases ^61^, as well as renal disorders ^62^. Distinguishing necrosis is crucial for accurate disease diagnosis and injury evaluation.

As depicted in **Figure 7a** for the liver, we constructed a logistic regression (LR) model using 226 real profiles, which were subsequently assessed on two test sets: one with real liver profiles and the other with matched synthetic profiles (see Methods section). Both test sets included 14 liver necrosis positives and 82 liver necrosis negatives. Results presented in **Table 1** indicate that the test set with synthetic profiles exhibited comparable performance to the test set with real profiles across all statistical measures.

**Figure 7:**
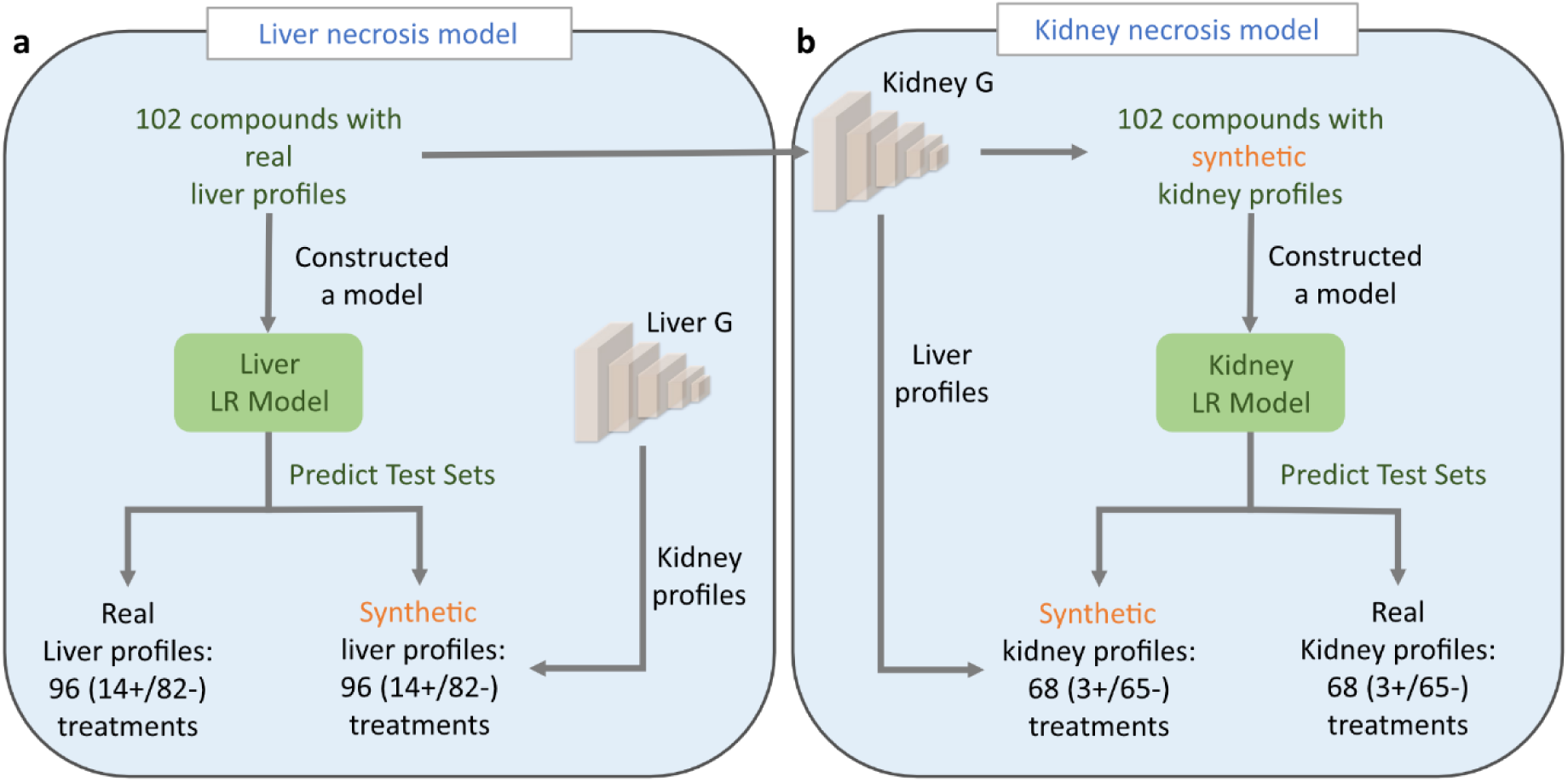
Framework of Necrosis Predictive Models. Liver Necrosis Predictive Model (**a**) - Developed using real liver profiles from 102 new compounds, the model was evaluated on two test sets: one with real liver profiles and the other with synthetic liver data generated by kidney generator. Kidney Necrosis Predictive Model (**b**) - Utilizing synthetic kidney profiles generated from the kidney generator with the liver profiles of 102 new compounds, the model was assessed on two test sets: real kidney data and synthetic kidney data generated from the liver generator.

**Table 1:**
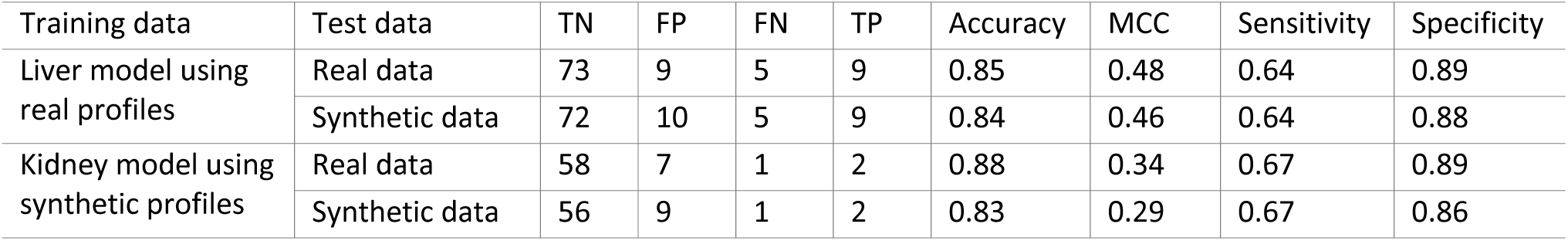
Performance of the liver and kidney necrosis predictive models on the test sets with synthetic or real transcriptomic data. TN: true negative; FP: false positive; FN: false negative; TP: true positive.

| Training data | Test data | TN | FP | FN | TP | Accuracy | MCC | Sensitivity | Specificity |
| --- | --- | --- | --- | --- | --- | --- | --- | --- | --- |
| Liver model using real profiles | Real data | 73 | 9 | 5 | 9 | 0.85 | 0.48 | 0.64 | 0.89 |
|  | Synthetic data | 72 | 10 | 5 | 9 | 0.84 | 0.46 | 0.64 | 0.88 |
| Kidney model using synthetic profiles | Real data | 58 | 7 | 1 | 2 | 0.88 | 0.34 | 0.67 | 0.89 |
|  | Synthetic data | 56 | 9 | 1 | 2 | 0.83 | 0.29 | 0.67 | 0.86 |

For the kidney, instead of utilizing real profiles as in the case of the liver, we developed an LR model using 52 synthetic profiles and tested it on two datasets: one with real kidney treatment profiles and the other with synthetic kidney treatment profiles (**Figure 7b**; see Methods section). Both datasets included 3 kidney necrosis positives and 65 kidney necrosis negatives. Similar performance was observed in the kidney, with the test set using synthetic profiles showing comparable performance to the test set with real profiles (**Table 1**).

## Discussion

Since its inception approximately 25 years ago, TGx has played a critical role in assessing chemical and drug safety. The majority of TGx studies have primarily concentrated on the liver due to (1) liver injury has been the top reasons for drugs failing in the clinical settings and (2) TGx experiments are costly to evaluate many organs. TransTox marks the initial AI effort in translating transcriptomic data between the liver and kidney under chemical treatment, providing a glimpse into the potential to extrapolate molecular expression to other organs based on the historical liver TGx studies. In a broader context, this investigation creates an opportunity to diminish the reliance on animal experimentation in future research focusing on organ toxicity. This is particularly applicable in cases where a wealth of liver data exists, thereby contributing to broader initiatives dedicated to reducing, refining, and replacing animal studies, as outlined in the 3Rs Principle ^63,64^.

TransTox was evaluated in multiple ways, most of which were focused on the results from the data that were not used in the training process (i.e., the test set and DrugMatrix data) and the scenarios that are relevant to the TGx applications (i.e., mechanistic interpretation and gene expression predictive models). The results demonstrated TransTox’s robust performance across all these tests.

DrugMatrix was applied to assess TransTox to utilize TGx data generated by other labs and predict compounds not seen in the training process. Selecting a validation dataset poses challenges due to diverse experimental designs, especially concerning TGx data, which exhibits significant variations not only in the choice of study design but also across different labs. It is crucial to choose an external dataset that allows an objective evaluation of TransTox results based on the model performance not due to the impact of differences in applied study designs. The study designs of DrugMatrix and TG-GATEs share commonalities but also have key differences, particularly regarding to dose range findings that affect dose definition (i.e., low, medium, and high). Fortunately, several compounds were tested in both TG-GATEs and DrugMatrix, serving as reference points to evaluate TransTox predictions. As depicted in **Figure 5**, the predicted results (correlation between synthetic and real data) aligned with experimental results (correlation between two datasets for common compounds), demonstrating its robustness for potential TGx applications.

Liver and kidney are distinct tissues with different mechanisms of action and varying responses to toxicants. Some compounds cause toxicity specifically to the kidneys, while others predominantly affect the liver. To demonstrate the reliability of TransTox predictions, we designed a study to evaluate its performance by focusing on organ-specific genes with a compound causing toxicity to a specific organ. In that way, agreement between prediction results and real data would indicate TransTox’s ability to handle organ-specific characteristics in translation. We analyzed thioacetamide-induced gene expression, as thioacetamide causes liver toxicity due to enterohepatic circulation with little impact on the kidneys. Our findings showed that the synthetic results closely matched real gene expression values, indicating that TransTox could capture the pattern of organ-specific gene expression, even with compounds that cause toxicity predominantly in specific organs.

One of the primary applications of TGx is to gain an understanding of underlying toxicity mechanisms through DEGs analysis and pathway extension. The concordance of DEGs across various gene expression platforms and labs has been extensively studied ^65^ and is largely dependent on chemical treatment and transcript abundance ^66^. The synthetic data produced by TransTox under specific treatment conditions can be likened to data obtained from the same experiment in a laboratory setting. In essence, the concordance of DEGs between synthetic and real data can be relatively evaluated by comparing it to that observed in experiments repeated multiple times. By following this logic, we selected a TGx experiment as reference points to evaluate the synthetic DEGs from TransTox. The experiment applied the similar study design as TG-GATEs including the microarray platforms to study two drugs (tolcapone and entacapone) with each experiment conditions involving 12 rats (4 groups of 3 rats to align with TG-GATEs design). We found that the percentage of agreement in DEGs between synthetic and real data is comparable to what is observed in experimental settings that were carried out in the same lab and by the same investigators but with different rats. A similar conclusion was drawn in pathway analysis, suggesting that TransTox could provide a synthetic approach for mechanistic studies in toxicology.

The other important TGx application is to develop predictive models for toxicity prediction. We evaluated the potential utility of synthetic profiles for predictive model development, addressing two primary questions: (1) whether an existing predictive model can be directly applied to synthetic profiles as “digital twin”, and (2) whether synthetic profiles could be used directly to develop predictive models. For questions 1, the liver model using real data achieved similar classification of necrosis in both real and synthetic profiles. For questions 2, the synthetic kidney predictive model achieved comparable performance to both real and synthetic profiles in distinguishing necrosis positives from negatives. These findings highlighted a possibility to use the synthetic data as “digital twin” to augment data size in predictive model development and evaluation in toxicology.

It is noted that TransTox was constructed using only 32 drugs which covered a limited chemistry space. In our previous work of assessing applicability domain in microarray-based genomic research ^67^, we noticed that gene expression data is less sensitive compared to QSARs approaches with respect to applicability domain. In this study, two external datasets were used to validate the model, including 9 drugs from TG-GATEs and 39 compounds from DrugMatrix, and the results are comparable to the experimental findings. With that said, our current evaluation in applicability domain was limited and the extensive application of TransTox to the diverse chemical universe required further studies. Therefore, the next iteration of TransTox will be focused on expanding the data size for both training, testing and validation. Due to computational constraints when this work was conducted, exhaustive testing of all hyperparameter combinations and model structures had not been extensively explored, which should also be carried out in the next iteration of the model development. Based on several GANs model developed for the TGx data including this one, we felt that the integration of existing biological knowledge, such as pathways, could be a valuable approach to improve the learning quality and, subsequently, improving the model’s overall performance.

The applications of generative methodologies in drug discovery and development are diverse^29,35^. Comparative analyses of these methods have been conducted. For example, Chen et al. compared synthetic data generated from five Differential Privacy (DP) generative models, including a GAN model, across three perspectives: downstream utility, statistical properties, and biological plausibility^68^. The study concluded that none of these methods accurately capture the biological characteristics of real datasets. Our GAN approach is based on a specific GAN architecture designed to translate information from one domain to another, with potential applications in translational science. However, a comprehensive comparative analysis of this method against similar AI methodologies has not been conducted in the field of toxicology. Given its potential utility in translational research, such an analysis would advance the field.

A significant challenge in toxicological research is the scarcity of available data samples and concerning animal usage, manifesting in three distinct scenarios. Firstly, regarding specific experimental technologies like transcriptomic methods, the absence of systematic evaluations across all organs impedes a systemic study of toxicity at the organism level. Secondly, regarding the animal use and 3Rs principle, in vitro experiments has been extensively evaluated in replacing in vivo studies, posing a challenge of in vitro-in vivo extrapolation (IVIVE). The third scenario involves multi-omics studies, where acquiring a complete set of omics data for an extensive collection of drugs or chemicals becomes especially challenging for comprehensive analysis. This study provided a glimpse of utilizing the advanced AI models for experimental data translation, offering a potential solution to extend the data diversity and to translate the data between experimental platforms.

## Methods

### Toxicogenomics (TGx) Dataset

The liver and kidney TGx datasets were obtained from the Open TG-GATEs ^51^, a large-scale TGx database including transcriptomic profiles and pathological findings from rat liver and kidney. The transcriptomic profiles were generated using Affymetrix GeneChip Rat Genome 230 2.0. Data from the Open TG-GATEs were downloaded from the website https://dbarchive.biosciencedbc.jp/en/open-tggates/download.html. Our study used the transcriptomic data from repeated dose experiments on rats, which employed three dose levels (low, middle, and high) and time-matched controls, along with four treatment durations (3, 7, 14, and 28 days).

The robust multi-array average (RMA) method ^69^ was applied to normalize the TG-GATEs data separately for both the liver and kidney. The expression values of the transcriptomic profiles were transformed to a log_2_ scale. To demonstrate the feasibility of organ translation using the TGx data, we focused on the rat S1500+ gene set. This gene set is considered important for toxicity by the Tox21 program and was identified through a hybrid approach comprising five sequential modules ^70^. This S1500+ gene set is specifically designed to represent biological diversity, address gene-gene co-expression relationships, and adequately represent known pathways of toxicity. The rat S1500+ gene set was downloaded from the website https://ntp.niehs.nih.gov/whatwestudy/tox21/s1500. Furthermore, we employed the “AnnotationDbi” package with the rat2302.db database to map the rat S1500+ genes to 3,475 microarray probes. From this point onward, the transcriptomic profiles refer to profiles containing the expression values of these 3,475 probes. The dataset had 41 compounds that had the paired transcriptomic data between liver and kidney. For each treatment conditions (combinations of compound, dose, and duration), 3 biological replicates (rats) were used; total of 616 treatment conditions comprised 1,844 profiles for each organ. Additionally, there were 102 compounds that had only liver profiles (no kidney data), with 3,595 liver profiles corresponding to 1,201 treatment conditions. For detailed information regarding the transcriptomic profiles and their associated pathological data, please refer to **Supplementary Data 3**.

### Training and Test Sets

To study organ translation between the liver and kidney, we employed drug-specific pairwise samples to train TransTox where each pairwise sample consisted of a liver profile and a kidney profile associated with a drug. For example, when considering acetaminophen as the drug, a liver profile under treatment condition A (e.g., dose=low and duration=7 days) was paired with a kidney profile under treatment condition B (dose=middle, duration=3 days), serving as a training sample for the kidney generator (using liver to generate kidney) in TransTox. Simultaneously, the kidney profile under treatment condition B was paired with the liver profile under treatment condition A, providing a training sample for the liver generator (using kidney to generate liver) in TransTox.

This study design was based on two well studied biomarkers in toxicology using gene expression profiles; these are early biomarker and sensitive biomarker. For a sensitive biomarker, its molecular expression at a low dose treatment could predict the phenotypic changes manifested at the high dose treatment condition^71–73^. For example, Dana et al. reported that changes in gene/protein expression generally correlated well with renal histopathological alterations and were often detected earlier or at lower doses than traditional clinical parameters like blood urea nitrogen and serum creatinine^71^. With respect to an early biomarker, its gene expression from a short-term experiment could predict toxicity which can only be found in a long-term study design^74–76^. For instance, Mark et al. reported that a novel multigene biomarker (i.e., signature) derived from short-term treatment could predict the likelihood of non-genotoxic chemicals inducing liver tumors in longer-term studies^75^. Therefore, we hypothesized that, within the same compound treatment, the toxicity relevant gene expression patterns present in every compound-specific treatment condition as such that these treatment samples can be paired to translate each other between different dosages and time points. Using these pairwise samples, the model was able to learn the complexity of compound-centric toxicity under different dose and time treatments.

To prevent information leakage, we divided the training and test sets based on compounds. This approach allocated 32 compounds to the training set and 9 compounds to the test set out of the total 41 compounds (**Table 2**). In addition to the three-dose treatments, we included control treatment profiles, enabling TransTox to thoroughly understand the underlying relationships between the liver and kidney. In this setup, a control treatment organ profile (e.g., liver) could only be paired with the control treatment profile of the same compound in the other organ (e.g., kidney). In total, there were 45,402 pairwise samples for each generator in the training set, while there were 11,250 pairwise samples in the test sets to evaluate the performance of both the liver and kidney generators.

**Table 2:**
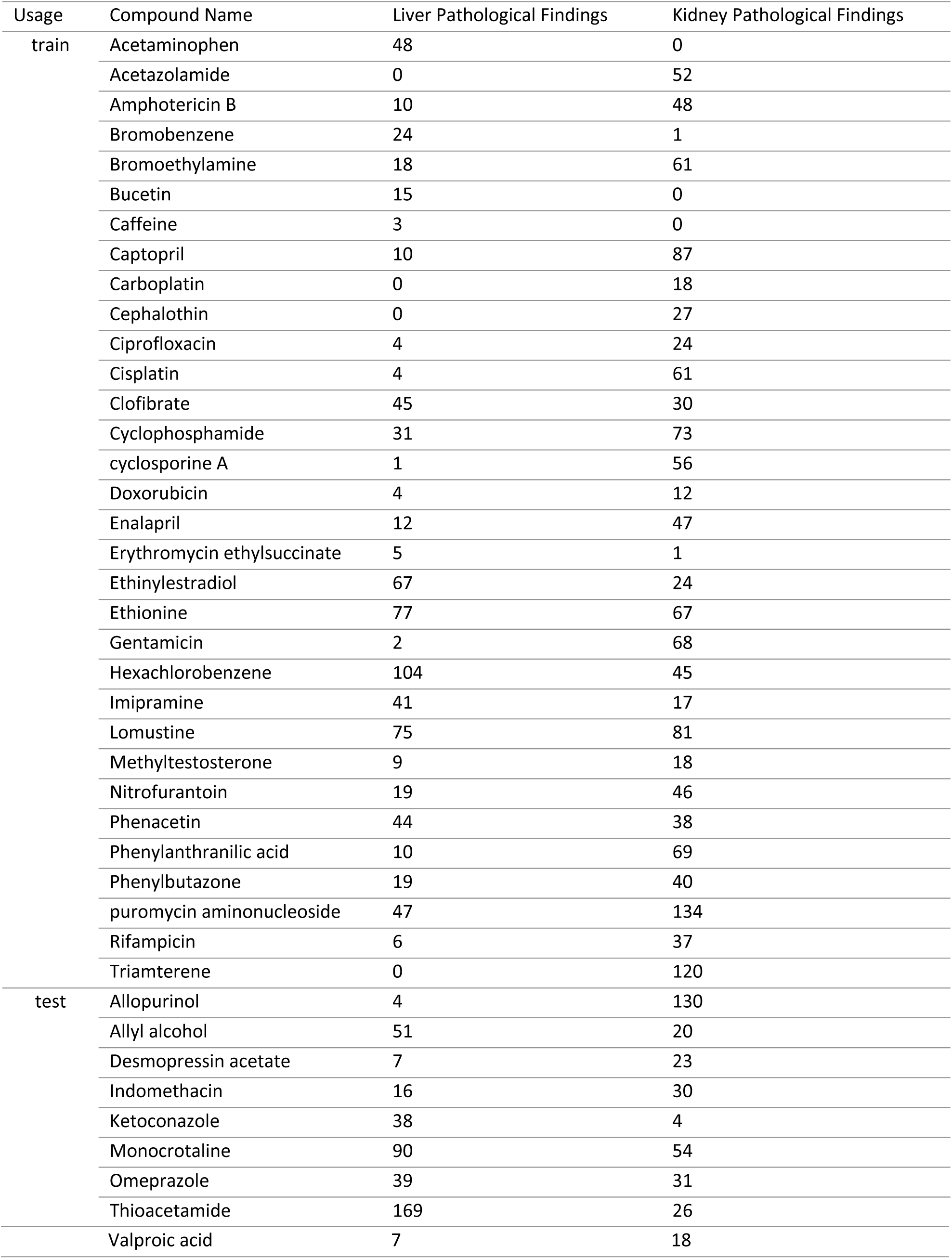
Compounds’ name and the number of pathological findings from TG-GATEs used in TransTox.

### TransTox Framework

TransTox was used to translate transcriptomic profiles under treatments from one organ (e.g., liver) to another (e.g., kidney), leveraging our previous model ^37^, which was developed using CycleGAN ^77^. Specifically, TransTox consisted of two pairs of generator and discriminator, one for liver and the other for kidney, interacting with each other as depicted in **Figure 1**. The liver Generator ((*G(liver)*) generated liver profiles from kidney profiles, while the Kidney Generator (*G(kidney)*) generated kidney profiles from liver profiles. Correspondingly, the Liver Discriminator (*D(liver)*) assessed the difference between synthetic and real data for the liver, while the Kidney Discriminator (*D(kidney)*) performed a similar function for the kidney. We developed a revised version of CycleGAN, implementing three key modifications to tailor it for organ translation using transcriptomic profiles. Firstly, we replaced the convolutional neural layers with dense layers to handle transcriptomic profiles as input data instead of images. Secondly, we enhanced the input layer by including additional labels to specify the origin and target attributes of the input profiles (e.g., organ, dose, time, etc.). Thirdly, we expanded the generator architecture with additional neurons to hold the labels of the target profile, thereby guiding the translation towards the desired direction.

Within the TransTox architecture, the generator input layer is designed to incorporate four essential components. The first component was the profile of the organ used for generating another organ—a 3,475-dimensional vector of transcriptomic values. The second component consisted of a 14-dimensional vector containing binary source label information, specifying details such as the organ, dose level, and time points. The third component was a 14-dimensional vector containing binary target label information. The fourth component involved a 3,475-dimensional Gaussian noise, sampled from a normal distribution, intended to enhance the model’s robustness. These four components were concatenated into a 6,978-dimensional vector and scaled to the range between 0 and 1. The generator was a fully connected neural network with five hidden layers, featuring 8,192, 7,168, 7,168, 4,096, and 4,096 nodes, respectively. Each hidden layer was followed by a dropout layer, except for the last one, with dropout rates of 0.8, 0.8, 0.8, and 0.4, respectively. The Leaky ReLU activation function, with an alpha value of 0.2, was applied across all hidden layers. The output layer was activated by a sigmoid activation function with 3,475 neurons, aligning with the dimensionality of features in the transcriptomic profiles.

The discriminator was a fully connected neural network consisting of four layers. The input layer comprised 3,475 neurons corresponding to the 3,475 probes, which received either the synthetic or real profile. Following this, there were two hidden layers with 256 and 64 neurons, respectively. The ReLU activation function was applied in each layer, followed by a dropout layer with a rate of 0.5. The output layer, consisting of a single neuron indicating whether the profile is real or synthetic, was activated by a sigmoid activation function. For optimization purposes, the discriminator model utilized Stochastic Gradient Descent (SGD) with a learning rate of 0.0001 and a momentum of 0.90.

### Assessing Synthetic Transcriptomic Profile from TransTox

In evaluating TransTox, we employed a comprehensive approach to assess its effectiveness in generating synthetic profiles in the test set. The first part of the assessment involved both visualization and quantitative analysis using three metrics. The UMAP technique ^52^ was selected for its ability to reduce high-dimensional data into a 2D representation, providing a visual overview of the distribution of synthetic and real profiles. Cosine similarity was applied to measure the likeness between synthetic and corresponding real profiles. The RMSE quantified the absolute differences between two profiles, offering insights into the accuracy of TransTox’s predictions. Additionally, the MAPE provided a percentage-wise measure of the errors between synthetic and real profiles.

### Gene-level comparison between the synthetic profiles and their corresponding real profiles for each organ and between organs

We evaluate gene-level variability within and between organs by comparing the difference between synthetic and real data using equations (1) and (2).

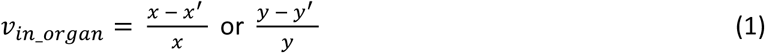

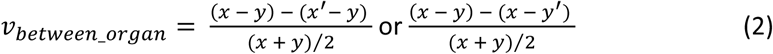

Here, *x* and *y* represent the real values, while *x’* and *y’* represents the synthetic gene expression data under treatment in liver and kidney, respectively. *v_in_organ_* denotes the variability of a gene between the synthetic data and real data in either liver or kidney, and *v_between_organ_* represents for the cross-organ variability for a gene between liver and kidney. The smaller absolute value of *v_in_organ_* and *v_between_organ_* indicates the difference between the real and synthetic values is small.

### External Validation of TransTox Predictions using DrugMatrix

DrugMatrix ^78^ was used as an external validation set to assess the accuracy of TransTox predictions for data generated from a different laboratory but with a similar study design. DrugMatrix is a large TGx database that includes transcriptomic profiles from primary tissues of male rats, including liver and kidney, profiled on the same Affymetrix Rat 230.2 platform as applied in TG-GATEs. The DrugMatrix data was downloaded from the Gene Expression Omnibus (https://www.ncbi.nlm.nih.gov/geo/, last accessed October 3, 2023) with accession numbers GSE57815 and GSE57811 for liver and kidney, respectively.

Two sequential criteria were applied to select compounds. Firstly, compounds with treatments lasting 3, 7, and 14 days were chosen to align with the TG-GATEs experimental time. Secondly, compounds needed to have both liver and kidney profiles, allowing for comparison between the generated profiles and the corresponding real profiles. In total, 39 compounds (**Table 3**) were selected, corresponding to 388 profiles (216 for liver and 172 for kidney). Five out of the 39 compounds were tested in both TG-GATEs and DrugMatrix, providing a matched reference for the analysis results. The remaining 34 compounds had incomplete profiles (i.e., either liver or kidney profiles, but not both) or no data in TG-GATEs, thus lacking a matched reference for the analysis results. Additionally, the treatment dose was converted to three levels (low, middle, and high) to maintain consistency with TG-GATEs dose information. The strategy for assigning doses was detailed in our previous study ^10^. The same microarray data preprocessing approach, RMA normalization, was employed to process the transcriptomic profiles from DrugMatrix. The processed liver profiles, including treatment information such as dose and time, were used as inputs for TransTox to generate the kidney profiles for the same compound, and vice versa. Subsequently, cosine similarity was applied to assess the similarity between the generated profiles from TransTox and the actual corresponding profiles from DrugMatrix.

**Table 3:** Compounds’ name and the number of pathological findings from DrugMatrix.

| In TG-GATEs | Compound Name | Liver Pathological Findings | Kidney Pathological Findings |
| --- | --- | --- | --- |
| No | 1-naphthyl isothiocyanate | 172 | 12 |
|  | Amiodarone | 0 | 3 |
|  | Atorvastatin | 41 | 4 |
|  | Azathioprine | 28 | 4 |
|  | Beta-estradiol | 19 | 3 |
|  | Bezafibrate | 82 | 3 |
|  | Bithionol | 19 | 0 |
|  | Carbon tetrachloride | 148 | 13 |
|  | Catechol | 18 | 3 |
|  | Cerivastatin | 49 | 0 |
|  | Cortisone | 9 | 0 |
|  | Daunorubicin | 14 | 0 |
|  | Dexamethasone | 14 | 0 |
|  | Diethylstilbestrol | 62 | 8 |
|  | Epirubicin | 25 | 3 |
|  | Erlotinib | 110 | 0 |
|  | Fenofibrate | 177 | 3 |
|  | Fluvastatin | 34 | 3 |
|  | Gefitinib | 139 | 0 |
|  | Gemfibrozil | 61 | 5 |
|  | Hydrocortisone | 14 | 3 |
|  | Ibuprofen | 37 | 22 |
|  | Imatinib | 16 | 3 |
|  | Lead (ii) acetate | 14 | 19 |
|  | Lead(iv) acetate | 16 | 12 |
|  | Leflunomide | 28 | 3 |
|  | Lovastatin | 39 | 3 |
|  | MIn-518 | 155 | 0 |
|  | N,n-dimethylformamide | 69 | 0 |
|  | N-nitrosodiethylamine | 63 | 0 |
|  | Naproxen | 51 | 17 |
|  | Nimesulide | 7 | 11 |
|  | Sodium arsenite | 41 | 0 |
|  | Sulindac | 64 | 24 |
| Yes | Allyl alcohol | 126 | 3 |
|  | Clofibrate | 87 | 3 |
|  | Doxorubicin | 13 | 0 |
|  | Gentamicin | 14 | 17 |
|  | Indomethacin | 81 | 26 |

### TransTox-Generated Transcriptomic Profiles in Elucidating Toxicity Mechanisms in Comparison with Real Experimental Settings

One important TGx application is to elucidate toxicity mechanisms. For that, we compared DEGs between synthetic profiles and their corresponding real profiles. Specifically, we calculated the average transcriptomic values of three biological replicates (rats) for a given treatment condition and compared them to that of the time-matched real control (i.e., three rats) to determine the log2-transformed fold change. For the real transcriptomic data, DEGs were defined as those with absolute log2-transformed fold changes greater than 1. The thresholds used to define DEGs for the synthetic profiles in the test set were based on the training set; the threshold was selected to ensure a similar average number of DEGs compared to the real data. In addition, we conducted KEGG pathway enrichment analyses separately for DEGs derived from synthetic profiles and real ones in each treatment condition using GSEApy ^79^. Enriched pathways with p-values < 0.05 were extracted for comparative analysis. This approach aimed to assess the concordance of DEGs and enriched pathways, providing insights into the potential of TransTox in elucidating toxicity mechanisms.

To evaluate the relevance of findings generated by TransTox (i.e., DEGs and pathways) to real experimental settings, we compared the TransTox results with a TGx experiment involving entacapone and tolcapone, which employed a study design similar to that of TG-GATEs ^80^. This dataset was curated from the US Food and Drug Administration’s (FDA’s) National Center for Toxicological Research (NCTR) ArrayTrack database ^81^. The experimental design, animal study execution, and the collection of the liver samples have been detailed in a prior study ^80^. In total, we collected 120 transcriptomic profiles sampled from male Sprague-Dawley rat liver tissue, including 10 treatments derived from the combination of two compounds (entacapone, tolcapone), four dose levels (control, low, middle, and high), and two dosing time periods (3- and 28-days). Specifically, the 3-day experiment used only control and high doses, resulting in 36 profiles (2 compounds x 12 rats + 1 control x 12 rats). In contrast, all treatment conditions were included in the 28-day study, producing 84 profiles (2 compounds x 12 rats x 3 doses + 1 control x 12 rats). The 120 transcriptomic profiles were processed with RMA normalization. Each treatment group comprised 12 biological replicates (i.e., 12 rats), which were further divided into four random subgroups, with three rats in each subgroup, consistent with the TG-GATEs study for a comparable comparison. Essentially, each subgroup represented an independent experiment, and the identical experiment was conducted four times by the same investigators in the same lab, concurrently, with only difference are the rats used. Subsequently, we analyzed the overlap in DEGs and pathways between any two subgroups, resulting in six comparisons for each treatment. These analyses serve as reference points for evaluating the performance of TransTox results within the context of the same experiment conducted multiple times with different rats.

### Necrosis Gene Expression Predictive Model Development

To explore the potential of TransTox in gene expression predictive model development, we focused on necrosis, a common pathological finding in both drug-induced liver and kidney injury. Necrosis also manifests in various acute and chronic liver diseases ^61^, as well as renal disorders ^62^. Distinguishing necrosis is fundamental for accurate disease diagnosis and injury assessment. A treatment is categorized as necrosis positive if at least one sample exhibits the necrosis pathological finding under that treatment. Conversely, a treatment is classified as necrosis negative if there are no pathological findings for all the rats in that treatment.

The model development focused on 102 compounds selected for specific reasons: (1) their liver profiles were available in TG-GATEs, and (2) their profiles were not used to develop or test TransTox. These 102 compounds consisted of 113 necrosis positives and 963 necrosis negatives in the liver, along with 26 necrosis positives and 664 necrosis negatives in the kidney.

In the liver, we first randomly selected 113 out of 963 treatment conditions without liver necrosis (negatives) to pair with the 113 treatment conditions with liver necrosis (positives) to construct a balanced training set. For each treatment condition, the average value across biological replicates were used. The liver necrosis model was developed using a logistic regression (LR) method. For the test set, all the treatment conditions with necrosis positive/negative data (real and synthetic) from the 9 different compounds were used to evaluate the liver LR model.

For the kidney, we were unable to develop a necrosis model based on real kidney profiles due to the absence of real data for modeling. Therefore, we generated synthetic kidney profiles using the kidney generator with real liver profiles and using these synthetic profiles to develop the kidney necrosis prediction model. Firstly, we randomly selected 26 out of 664 treatment conditions without kidney necrosis (negatives). Secondly, these were combined with the 26 treatment conditions with kidney necrosis (positives) to create a balanced training set. Thirdly, the average value across biological replicates was calculated for each treatment condition and further utilized to train the kidney LR model for predicting kidney necrosis. Finally, the model was tested on the treatment conditions with necrosis positive/negative data (real and synthetic) from the 9 different compounds.

To quantitatively assess the models’ performance on two different test datasets, we utilized four performance metrics: Matthews correlation coefficient (MCC), accuracy, sensitivity, and specificity. These measurements are calculated using equations (3) through (6).

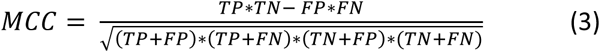

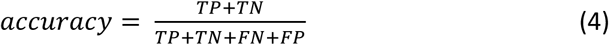

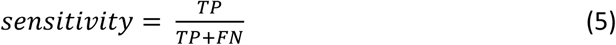

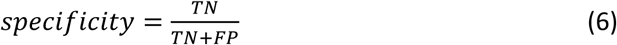

In these formulas, TP, TN, FP, and FN represent true positive, true negative, false positive, and false negative, respectively. Further elaboration on these definitions can be found elsewhere ^82^.

## Data Availability

The datasets used to develop and evaluate TransTox are collected from the Open TG-GATEs database, available for downloaded at https://dbarchive.biosciencedbc.jp/en/open-tggates/download.html. The DrugMatrix data was downloaded from the Gene Expression Omnibus at https://www.ncbi.nlm.nih.gov/geo/, with accession numbers GSE57815 and GSE57811 for liver and kidney, respectively.

## Code Availability

TransTox was developed using tensorflow-GPU version 2.2.0 with open-source Python (version 3.7.11). The source code is available at https://github.com/TingLi2016/TransTox.

## Acknowledgements

We would like to express our sincere appreciation to Dr. Tao Han for providing the microarray data for entacapone and tolcapone. We are also grateful to Dr. Dongying Li for providing the biological insights included in our response letter. Both are from the FDA National Center for Toxicological Research. The views presented in this article do not necessarily reflect those of the U.S. Food and Drug Administration. Any mention of commercial products is for clarification and is not intended as an endorsement. No funding was granted for the study.

## Author Contributions

W.T. conceived and designed the study. T.L. performed the model development and computational analysis. X.C. provided data and interpretation of the results. T.L. wrote the first draft of the manuscript. W.T. revised the manuscript. All authors edited and approved the final manuscript.

## Reference

1 Su, Z. et al. An investigation of biomarkers derived from legacy microarray data for their utility in the RNA-seq era. Genome biology 15, 1–25 (2014).

2 Huang, J. et al. Genomic indicators in the blood predict drug-induced liver injury. The pharmacogenomics journal 10, 267–277 (2010).

3 Gibney, E. & Nolan, C. Epigenetics and gene expression. Heredity 105, 4–13 (2010).

4 Romero, I. G., Ruvinsky, I. & Gilad, Y. Comparative studies of gene expression and the evolution of gene regulation. Nature Reviews Genetics 13, 505–516 (2012).

5 Bar-Joseph, Z., Gitter, A. & Simon, I. Studying and modelling dynamic biological processes using time-series gene expression data. Nature Reviews Genetics 13, 552–564 (2012).

6 Hill, M. S., Vande Zande, P. & Wittkopp, P. J. Molecular and evolutionary processes generating variation in gene expression. Nature Reviews Genetics 22, 203–215 (2021).

7 De Nadal, E., Ammerer, G. & Posas, F. Controlling gene expression in response to stress. Nature Reviews Genetics 12, 833–845 (2011).

8 Nilsen, T. W. Mechanisms of microRNA-mediated gene regulation in animal cells. TRENDS in Genetics 23, 243–249 (2007).

9 Gräff, J., Kim, D., Dobbin, M. M. & Tsai, L.-H. Epigenetic regulation of gene expression in physiological and pathological brain processes. Physiological reviews 91, 603–649 (2011).

10 Chen, X., Roberts, R., Tong, W. & Liu, Z. Tox-GAN: an artificial intelligence approach alternative to animal studies—a case study with toxicogenomics. Toxicological Sciences 186, 242–259 (2022).

11 Alexander-Dann, B. et al. Developments in toxicogenomics: understanding and predicting compound-induced toxicity from gene expression data. Molecular omics 14, 218–236 (2018).

12 Wu, Y. & Wang, G. Machine learning based toxicity prediction: from chemical structural description to transcriptome analysis. International journal of molecular sciences 19, 2358 (2018).

13 Seal, S. et al. Integrating cell morphology with gene expression and chemical structure to aid mitochondrial toxicity detection. Communications Biology 5, 858 (2022).

14 Li, T., Tong, W., Roberts, R., Liu, Z. & Thakkar, S. Deep learning on high-throughput transcriptomics to predict drug-induced liver injury. Frontiers in bioengineering and biotechnology 8, 562677 (2020).

15 Waring, J. F. et al. Clustering of hepatotoxins based on mechanism of toxicity using gene expression profiles. Toxicology and applied pharmacology 175, 28–42 (2001).

16 Tan, Y., Shi, L., Tong, W., Hwang, G. G. & Wang, C. Multi-class tumor classification by discriminant partial least squares using microarray gene expression data and assessment of classification models. Computational Biology and Chemistry 28, 235–243 (2004).

17 Yang, W. et al. Identification of genes and pathways involved in kidney renal clear cell carcinoma. BMC bioinformatics 15, 1–10 (2014).

18 Fang, H. et al. Gene expression profile exploration of a large dataset on chronic fatigue syndrome. (2006).

19 San Segundo-Val, I. & Sanz-Lozano, C. S. Introduction to the gene expression analysis. Molecular genetics of asthma, 29–43 (2016).

20 Pugazhendhi, A., Edison, T. N. J. I., Velmurugan, B. K., Jacob, J. A. & Karuppusamy, I. Toxicity of Doxorubicin (Dox) to different experimental organ systems. Life sciences 200, 26–30 (2018).

21 Andjelkovic, M. et al. Toxic effect of acute cadmium and lead exposure in rat blood, liver, and kidney. International journal of environmental research and public health 16, 274 (2019).

22 Renu, K., Pureti, L. P., Vellingiri, B. & Valsala Gopalakrishnan, A. Toxic effects and molecular mechanism of doxorubicin on different organs–an update. Toxin Reviews 41, 650–674 (2022).

23 Yang, Y. et al. Toxicity assessment of nanoparticles in various systems and organs. Nanotechnology Reviews 6, 279–289 (2017).

24 Denny, K. H. in A comprehensive guide to toxicology in nonclinical drug development 149–171 (Elsevier, 2024).

25 Oleaga, C. et al. Multi-Organ toxicity demonstration in a functional human in vitro system composed of four organs. Scientific reports 6, 20030 (2016).

26 Esch, M. B. et al. How multi-organ microdevices can help foster drug development. Advanced drug delivery reviews 69, 158–169 (2014).

27 Esch, M., King, T. & Shuler, M. The role of body-on-a-chip devices in drug and toxicity studies. Annual review of biomedical engineering 13, 55–72 (2011).

28 Raies, A. B. & Bajic, V. B. In silico toxicology: computational methods for the prediction of chemical toxicity. Wiley Interdisciplinary Reviews: Computational Molecular Science 6, 147–172 (2016).

29 Méndez-Lucio, O., Baillif, B., Clevert, D.-A., Rouquié, D. & Wichard, J. De novo generation of hit-like molecules from gene expression signatures using artificial intelligence. Nature communications 11, 10 (2020).

30 Kim, H., Kim, Y., Lee, C.-Y., Kim, D.-G. & Cheon, M. Investigation of early molecular alterations in tauopathy with generative adversarial networks. Scientific Reports 13, 732 (2023).

31 Tong, X. et al. Generative models for de novo drug design. Journal of Medicinal Chemistry 64, 14011–14027 (2021).

32 Macedo, B., Ribeiro Vaz, I. & Taveira Gomes, T. MedGAN: optimized generative adversarial network with graph convolutional networks for novel molecule design. Scientific Reports 14, 1212 (2024).

33 Green, A. J. et al. Leveraging high-throughput screening data, deep neural networks, and conditional generative adversarial networks to advance predictive toxicology. PLOS Computational Biology 17, e1009135 (2021).

34 Umarov, R., Li, Y. & Arner, E. DeepCellState: An autoencoder-based framework for predicting cell type specific transcriptional states induced by drug treatment. PLoS Computational Biology 17, e1009465 (2021).

35 Chen, X., Roberts, R., Liu, Z. & Tong, W. A generative adversarial network model alternative to animal studies for clinical pathology assessment. Nature Communications 14, 7141 (2023).

36 Ge, Q. et al. Conditional generative Adversarial networks for individualized treatment effect estimation and treatment selection. Frontiers in genetics 11, 585804 (2020).

37 Li, T., Roberts, R., Liu, Z. & Tong, W. TransOrGAN: An Artificial Intelligence Mapping of Rat Transcriptomic Profiles between Organs, Ages, and Sexes. Chemical Research in Toxicology (2023).

38 Burcham, P. C. & Burcham, P. C. Target-organ toxicity: liver and kidney. An Introduction to Toxicology, 151–187 (2014).

39 Li, X., Hassoun, H. T., Santora, R. & Rabb, H. Organ crosstalk: the role of the kidney. Current opinion in critical care 15, 481–487 (2009).

40 Serteser, M. et al. Changes in hepatic TNF-α levels, antioxidant status, and oxidation products after renal ischemia/reperfusion injury in mice. Journal of surgical research 107, 234–240 (2002).

41 Capalbo, O., Giuliani, S., Ferrero-Fernández, A., Casciato, P. & Musso, C. G. Kidney–liver pathophysiological crosstalk: Its characteristics and importance. International Urology and Nephrology 51, 2203–2207 (2019).

42 Yap, S. C., Lee, H. T. & Warner, D. S. Acute kidney injury and extrarenal organ dysfunction: new concepts and experimental evidence. The Journal of the American Society of Anesthesiologists 116, 1139–1148 (2012).

43 Wadei, H. M. in Seminars in respiratory and critical care medicine. 55-69 (Thieme Medical Publishers).

44 Bonavia, A. & Stiles, N. Renohepatic crosstalk: a review of the effects of acute kidney injury on the liver. Nephrology Dialysis Transplantation 37, 1218–1228 (2022).

45 Golab, F. et al. Ischemic and non-ischemic acute kidney injury cause hepatic damage. Kidney international 75, 783–792 (2009).

46 Moore, J. K., Love, E., Craig, D. G., Hayes, P. C. & Simpson, K. J. Acute kidney injury in acute liver failure: a review. Expert review of gastroenterology & hepatology 7, 701–712 (2013).

47 Zhao, M. et al. Cytochrome P450 enzymes and drug metabolism in humans. International journal of molecular sciences 22, 12808 (2021).

48 George, B., You, D., Joy, M. S. & Aleksunes, L. M. Xenobiotic transporters and kidney injury. Advanced drug delivery reviews 116, 73–91 (2017).

49 Jetter, A. & Kullak-Ublick, G. A. Drugs and hepatic transporters: A review. Pharmacological research 154, 104234 (2020).

50 Rosenthal, S. B., Bush, K. T. & Nigam, S. K. A network of SLC and ABC transporter and DME genes involved in remote sensing and signaling in the gut-liver-kidney axis. Scientific reports 9, 11879 (2019).

51 Igarashi, Y. et al. Open TG-GATEs: a large-scale toxicogenomics database. Nucleic acids research 43, D921–D927 (2015).

52 McInnes, L., Healy, J. & Melville, J. Umap: Uniform manifold approximation and projection for dimension reduction. arXiv preprint arXiv:1802.03426 (2018).

53 Chiang, J. Y. & Ferrell, J. M. Up to date on cholesterol 7 alpha-hydroxylase (CYP7A1) in bile acid synthesis. Liver research 4, 47–63 (2020).

54 Pan, Y., Chen, H. & Yu, J. Biomarkers in hepatocellular carcinoma: current status and future perspectives. Biomedicines 8, 576 (2020).

55 Kwon, T.-H., Frøkiaer, J., Knepper, M. A. & Nielsen, S. Reduced AQP1,-2, and-3 levels in kidneys of rats with CRF induced by surgical reduction in renal mass. American Journal of Physiology-Renal Physiology 275, F724–F741 (1998).

56 Nielsen, S. et al. Aquaporins in the kidney: from molecules to medicine. Physiological reviews 82, 205–244 (2002).

57 Narlis, M., Grote, D., Gaitan, Y., Boualia, S. K. & Bouchard, M. Pax2 and pax8 regulate branching morphogenesis and nephron differentiation in the developing kidney. Journal of the American Society of Nephrology 18, 1121–1129 (2007).

58 Crespo Yanguas, S., et al. Experimental models of liver fibrosis. Archives of toxicology 90, 1025–1048 (2016).

59 Schyman, P. et al. Assessing chemical-induced liver injury in vivo from in vitro gene expression data in the rat: The case of thioacetamide toxicity. Frontiers in Genetics 10, 1233 (2019).

60 Schyman, P. et al. Identification of the toxicity pathways associated with thioacetamide-induced injuries in rat liver and kidney. Frontiers in Pharmacology 9, 1272 (2018).

61 Krishna, M. Patterns of necrosis in liver disease. Clinical liver disease 10, 53 (2017).

62 Belavgeni, A., Meyer, C., Stumpf, J., Hugo, C. & Linkermann, A. Ferroptosis and necroptosis in the kidney. Cell chemical biology 27, 448–462 (2020).

63 Clark, J. M. The 3Rs in research: a contemporary approach to replacement, reduction and refinement. British Journal of Nutrition 120, S1–S7 (2018).

64 House of Lords (2002) Report of the Select Committee on animals in scientific procedures., <https://publications.parliament.uk/pa/ld200102/ldselect/ldanimal/150/150.pdf> (

65 Shi, L. et al. The MicroArray Quality Control (MAQC) project shows inter-and intraplatform reproducibility of gene expression measurements. Nature biotechnology 24, 1151–1161 (2006).

66 Wang, C. et al. The concordance between RNA-seq and microarray data depends on chemical treatment and transcript abundance. Nature biotechnology 32, 926–932 (2014).

67 Shao, L., Wu, L., Fang, H., Tong, W. & Fan, X. Does applicability domain exist in microarray-based genomic research? PLoS One 5, e11055 (2010).

68 Chen, D., et al. Towards biologically plausible and private gene expression data generation. arXiv preprint arXiv:2402.04912 (2024).

69 Irizarry, R. A. et al. Exploration, normalization, and summaries of high density oligonucleotide array probe level data. Biostatistics 4, 249–264 (2003).

70 Mav, D. et al. A hybrid gene selection approach to create the S1500+ targeted gene sets for use in high-throughput transcriptomics. PloS one 13, e0191105 (2018).

71 Hoffmann, D. et al. Performance of novel kidney biomarkers in preclinical toxicity studies. Toxicological Sciences 116, 8–22 (2010).

72 Wetmore, B. A. et al. Quantitative analyses and transcriptomic profiling of circulating messenger RNAs as biomarkers of rat liver injury. Hepatology 51, 2127–2139 (2010).

73 Anadón, A., Castellano, V. & Martínez-Larrañaga, M. R. in Biomarkers in toxicology 593–607 (Elsevier, 2014).

74 Mina, S. G. et al. Assessment of drug-induced toxicity biomarkers in the brain microphysiological system (MPS) using targeted and untargeted molecular profiling. Frontiers in big Data 2, 23 (2019).

75 Fielden, M. R., Brennan, R. & Gollub, J. A gene expression biomarker provides early prediction and mechanistic assessment of hepatic tumor induction by nongenotoxic chemicals. Toxicological sciences 99, 90–100 (2007).

76 Corton, J. C., Hill III, T., Sutherland, J. J., Stevens, J. L. & Rooney, J. A set of six gene expression biomarkers identify rat liver tumorigens in short-term assays. Toxicological Sciences 177, 11–26 (2020).

77 Zhu, J.-Y., Park, T., Isola, P. & Efros, A. A. in Proceedings of the IEEE international conference on computer vision. 2223–2232.

78 Ganter, B., Snyder, R. D., Halbert, D. N. & Lee, M. D. Toxicogenomics in drug discovery and development: mechanistic analysis of compound/class-dependent effects using the DrugMatrix® database. (2006).

79 Fang, Z., Liu, X. & Peltz, G. GSEApy: a comprehensive package for performing gene set enrichment analysis in Python. Bioinformatics 39, btac757 (2023).

80 McBurney, R. N. et al. The liver toxicity biomarker study: phase I design and preliminary results. Toxicologic Pathology 37, 52–64 (2009).

81 Tong, W. et al. ArrayTrack--supporting toxicogenomic research at the US Food and Drug Administration National Center for Toxicological Research. Environmental health perspectives 111, 1819–1826 (2003).

82 Li, T., Liu, Z., Thakkar, S., Roberts, R. & Tong, W. DeepAmes: A deep learning-powered Ames test predictive model with potential for regulatory application. Regulatory Toxicology and Pharmacology 144, 105486 (2023).

