## Supplementary figures and images for "Bridging Organ Transcriptomics for Advancing Multiple Organ Toxicity Assessment with a Generative AI Approach"

### Supplementary Figure 1

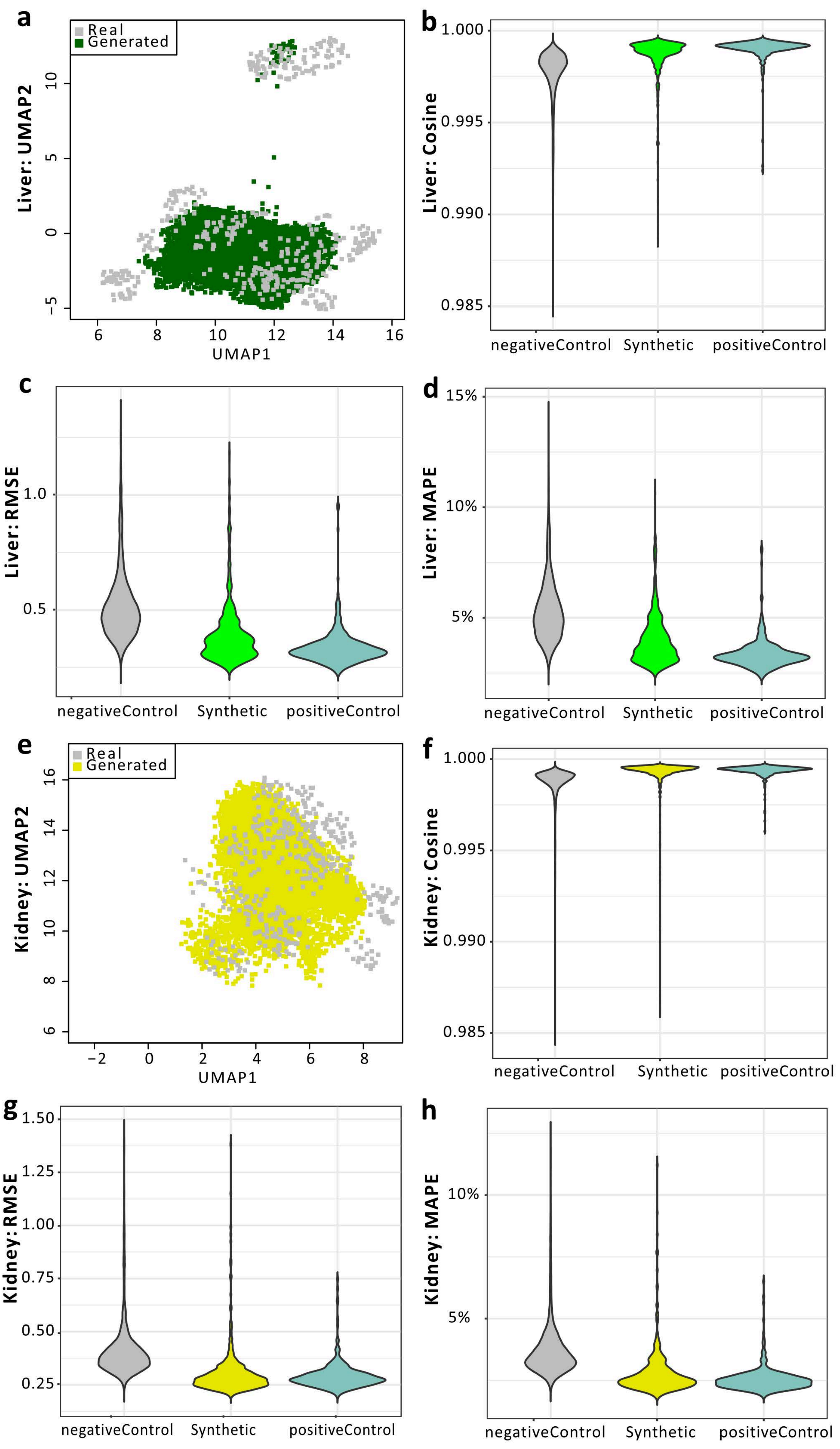
